# Top-down structuring of freshwater bacterial communities by mixotrophic and heterotrophic protists

**DOI:** 10.1101/2022.08.12.503741

**Authors:** Marina Ivanković, Robert Ptacnik, Mia Maria Bengtsson

## Abstract

Mixotrophic and heterotrophic protists hold a key position in aquatic microbial food webs. They account for the bulk of bacterivory in pelagic systems. However, the potential structuring effect of heterotrophic and especially mixotrophic protists on bacterial communities is far from clear. We conducted standardized short-term grazing experiments, to test for the overall impact on bacterial community structure and possible prey preferences of protist taxa with different phagotrophic nutritional modes. The used protist taxa covered a range from more phototrophic towards heterotrophic strategies, with obligate mixotrophic taxa employing phototrophy as the dominant strategy, towards facultative mixotrophic taxa relying more on phagotrophy, with the end of this gradient being covered by a phagoheterotroph lacking phototrophic capacity. Bacterioplankton from different lake systems was enriched and used to represent semi-natural bacterial assemblages that served as prey communities. Our study showed that similarities in protistan nutritional modes were reflected in similarities in their overall impact on bacterial communities. The impact intensity increased towards clear phagotrophic strategies, with mixotrophs in the phototrophic end of the gradient rather having a stabilizing impact on bacterial communities. Obligate mixotrophs grazed diverse prey in small amounts, while towards strict heterotrophic nutrition we observed an ingestion increase and selective depletion of bacteria with potential high growth rates. Future global change scenarios supporting the domination of obligate mixotrophic bacterivores might promote higher bacterial diversity in the illuminated zone of lakes.

## Introduction

Bacteria hold a key position in essentially all biogeochemical cycles within aquatic ecosystems, playing an important role as degraders and remineralizers of particulate organic matter, while simultaneously forming the base of most pelagic food webs either as primary or secondary producers (1–3). The interactions of bacteria with other organisms determine the rate and efficiency of these processes. Yet, most studies so far mainly focused only on bottom-up related bacterial interactions, while top-down interactions such as predation and viral lysis are far less studied und understood. It is well known that predation upon bacteria by protists has a crucial position within the complex network of bacterial interactions (1,4). Protist bacterivory not only controls the abundance and biomass of bacterial populations, it also structures bacterial community composition (5). Consequently, protists play a pivotal role as a factor influencing the overall ecological function of bacterial communities, influencing energy fluxes, nutrient transfer and recycling within the microbial loop (6,7).

Nano-sized mixotrophic and heterotrophic protists are recognized as main consumers of bacteria in the pelagial, thereby themselves holding a key position in aquatic food webs and overall pelagic ecosystem functioning (5,8,9). The term mixotrophy referrers here to protists combining phototrophy with phagotrophic uptake of prey. Existing studies mostly focused on heterotrophic taxa as bacterivores (5), and references inside), while mixotrophic predation, its functional variation and implications for bacterial community structuring were rarely addressed. This is surprising, given the fact that mixotrophic bacterivory is a common trait in freshwater and marine plankton, often of at least equal importance in the removal of various picoplanktonic cells as predation by heterotrophic bacterivores (8,10–12).

The variety of empirical observations and described mixotrophic taxa are suggesting that the mixotrophic nutrition is covering a gradient from obligate heterotrophy to phototrophic nutrition (13). Different mixotrophic taxa can cover a wider or narrower range between these two nutritional strategies, by changing their phototrophic:phagotrophic lifestyle ratio in relation to the existing environmental conditions (9,14,15). In general, the dual strategy of mixotrophic protists enables them to cover nutrients and energy demands from alternative sources. They can utilize essential nutrients from ingested prey and from the surrounding water, while covering necessary energy and carbon demands over photosynthesis and prey ingestion (13). Simple functional classifications often reflect the main nutritional strategy of mixotrophic taxa, resulting with two main groups, obligate and facultative mixotrophs (9,16,17). Growth of facultative mixotrophs is primarily heterotrophic, mainly depends on prey supply, with light and inorganic nutrients often serving as minor resources (18–21). Conversely, obligate mixotrophic taxa are primarily phototrophic, mainly rely on photosynthesis, and ingest lesser amounts of bacteria mainly depending on the availability of dissolved nutrients (21–24). Today, the quantitative dependence on bacterial prey of these two mixotrophic groups is relatively well known, in contrast to the influence they have on bacterial community composition.

Feeding preferences of consumers are generally expected to reflect their nutritional needs, related to the consumer cellular composition and physiology (25–28). While certain prey may be preferred due its nutritional quality, other factors will influence the outcome of bacteria-protist interactions. First, the probability that protist and bacteria encounter each other, with motility and abundance of both groups being important (29–31). Second, grazing-resistant strategies of bacteria play a crucial role, whereby size, morphological traits or biochemical protection via different substances can act repelling or harmful on the protist (see reviews by (32–34). Finally, the growth rate of the bacterial prey may determine their ability to compensate for grazing loss and thereby persist in the environment (31). Whereas these factors set the fundament for the complex interactions happening between protist and bacteria, still little is known how protist bacterivory contributes to bacterial community structure.

We conducted standardized short-term grazing experiments, to examine the relation of different trophic nutritional modes in protist taxa to their overall impact on bacterial communities and possible prey selectivity patterns. The employed protist taxa covered a range from more phototrophic towards heterotrophic strategies, with obligate mixotrophic taxa employing phototrophy as the dominant strategy, towards facultative mixotrophic taxa relying more on phagotrophy, and a phagoheterotroph lacking phototrophic capacity. We collected natural plankton communities from different lakes and enriched them to create a variation of semi-natural bacterial communities serving as prey. The objective of the study was to assess the influence of the selected protists on diverse bacterial communities. We hypothesized that the impact of protists on bacterial communities follows the gradient from phototrophy towards phagotrophy, with protists taxa being closer within the continuum having a more similar impact on bacterial communities than taxa being more distant.

## Material and methods

### Experimental organisms

Four bacterivorous protist taxa were employed as bacterivores, three mixotrophic flagellates: *Uroglenopsis americana, Ochromonas* c.f. *perlata* and *Poteriochromonas malhamensis*, and one heterotrophic *Spumella*-like flagellate. The protists are hereinafter referred to with their genus name. These flagellates are taxonomically closely related and similar in their morphology, but represent different trophic modes, from primarily phototrophic obligate mixotrophs (*Uroglenopsis, Ochromonas*) to primarily phagotrophic facultative mixotrophs (*Poterioochromonas*) and a phagoheterotroph lacking phototrophic capacity (*Spumella*). For a detailed description addressing the protists see Table 1. The prey communities originate from seven different natural lakes: Lunzer See, Mittersee, Obersee, Erlaufsee, Hubertussee, Klostersee and Chiemsee (see Table S1 for Lake description). In order to test for general trends in terms of protist-bacteria interactions, the chosen lakes represent a range of different potential habitats for the consumers employed in this study.

**Table 1.**
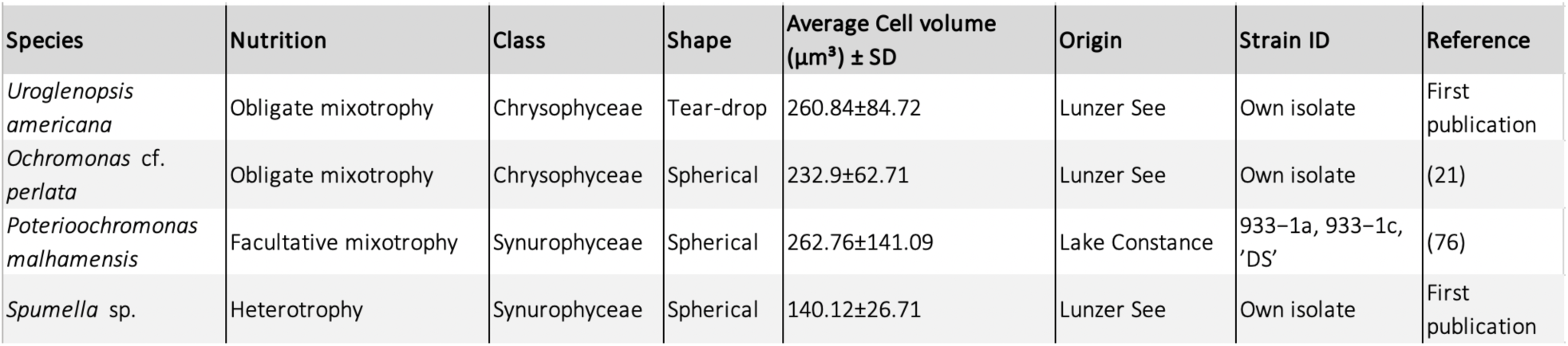
Characteristics of protist taxa used as bacterial consumers.

| Species | Nutrition | Class | Shape | Average Cell volume ($\mu\text{m}^3$) ± SD | Origin | Strain ID | Reference |
| --- | --- | --- | --- | --- | --- | --- | --- |
| *Uroglenopsis americana* | Obligate mixotrophy | Chrysophyceae | Tear-drop | 260.84±84.72 | Lunzer See | Own isolate | First publication |
| *Ochromonas* cf. *perlata* | Obligate mixotrophy | Chrysophyceae | Spherical | 232.9±62.71 | Lunzer See | Own isolate | (21) |
| *Poterioochromonas malhamensis* | Facultative mixotrophy | Synurophyceae | Spherical | 262.76±141.09 | Lake Constance | 933–1a, 933–1c, 'DS' | (76) |
| *Spumella* sp. | Heterotrophy | Synurophyceae | Spherical | 140.12±26.71 | Lunzer See | Own isolate | First publication |

### Cultivation and preparation of prey bacteria and protist prior to the experiment

All organisms were pre-cultivated in autoclaved glass bottles placed in a walk-in environmental chamber, at a constant temperature of 18°C and a light:dark cycle of 16:8h, with a smooth transition between light and dark phases. The light source consisted of three different LEDs, mimicking natural PAR, supplying non-limiting irradiance of ca. 80 µmol m^−2^ s^−1^. The handling of the lake prey communities, as well of the protist cultures prior and during the experiment was done under a laminar flow hood under sterile conditions.

Bacterial communities for prey preparation were sampled from the epilimnion of every lake, by taking a 10 L water sample with a clean container rinsed with lake water. The water samples where immediately transported to the laboratory under dark and cooled conditions and filtered by gentle vacuum filtration through 0.8 μm polycarbonate membrane filters to exclude protists and any larger organisms. The filtrate was then pelleted (3000 G, 10 min, RT) and resuspended in >10 weeks aged 0.2 µm sterile filtered Lunzer See water. Organic carbon and inorganic phosphorus were added according to the Redfield ratio (Glucose, 83 µmol C L^-1^, Monopotassium phosphate, 0.79 µmol P L^-1^) to support bacterial growth. No nitrogen source was added, as nitrogen is a non-limiting nutrient in the Lunzer See (35). The prey communities were kept in exponential growth for 48h on the enriched medium until further processing.

The protist cultures where pre-grown on medium based on sterile filtered lake water (same as used for bacteria) with a 5% final addition of modified WEES medium (WEES Medium Recipe v.03.2007 without soil extract (36)). Protists were kept in exponential growth prior every experiment. In order to separate the protist cells from the culture medium, all protists, except *Uroglenopsis*, were pelleted via gentle centrifugation (3000 G, 10 min, RT), and subsequently resuspended in sterile 0.2 µm filtered Lunzer See water. *Uroglenopsis* is forming colonies which are very sensitive to mechanical stress and was therefore concentrated via light attraction, and carefully resuspended via pipetting. After resuspension the cultures were kept undisturbed for 24h to adapt to the sterile lake water and to recover from any stress induced by centrifugation or pipetting (the used recovery time turned out to prevent negative effects on flagellate growth and ingestion, tested in preliminary experiments). Due to low densities during the pre-cultivation, *Ochromonas* was not employed in the Lunzer See and Obersee experiments. The protist cultures used in this study were not axenic, i.e. they contained heterotrophic bacteria.

### Experimental setup, incubation and sample processing

The natural lake communities were used to create a number of semi-natural, diverse bacterial communities, serving as prey. After the pre-cultivation and recovery phase, the prey community from one lake was separately inoculated with each protist, being further on referred as ‘Protist treatments’, plus with one control treatment containing only the bacterial prey (‘Control’). Since the protist cultures were not axenic, bacteria were introduced along with the protist culture, specific for every protist treatment, but not present in the control, further on called ‘Background bacteria’. Per lake and treatment, a volume of 2 L was prepared, gently shaken and split into four 500 ml autoclaved glass bottles. One bottle was immediately harvested (‘Start’ of experiment), the remaining three bottles where incubated for 48 h and harvested (‘End’ of experiment). The experimental incubations were carried out under identical conditions as described above for the pre-cultivation phase. The volume of every bottle was used for flow cytometry, microscopic and molecular analyses. Samples of 20 ml for flow cytometry and fluorescence microscopy were fixed with a mixture of 0.2 µm filtered paraformaldehyde and glutaraldehyde with a final concentration of 0.01% and 0.1%, respectively, in the sample, and subsequently stored at 4°C until analysis (within 24h after fixation). A volume of 400 ml was filtered on polyethersulfone 0.2 μm membrane filters and kept frozen at -80°C until DNA extraction.

### Prey and protist enumeration, growth and grazing parameters estimation

Cell numbers were estimated via flow cytometry using a Beckman Coulter CytoFLEX flow cytometer (Beckman Coulter GmbH, Krefeld, Germany), data acquisition and analysis were carried out using the CytExpert Software 2.3. To count heterotrophic organisms, subsamples were stained with a SYTO 13 (Invitrogen, final concentration 0.5 μM). After stain addition, the sample was kept 15 minutes at RT in the dark and subsequently analyzed via flow cytometry as described below. Stained bacteria and *Spumella* cells were gated on a dot plot of blue absorption light and green fluorescence versus the forward scatter. Using unstained samples, the mixotrophs were gated by their Chl-*a* auto-fluorescence signal by using a scatter plot of blue excitation light and red fluorescence light versus the forward scatter. The thresholds were set to minimize background noise, and the organisms were gated manually. In addition, fluorescence and light microscopy were used to validate flow cytometer templates, as well to check that no contamination by the protist organisms between treatments happened. Net growth rates of bacteria and protists in the incubation bottles were calculated for each treatment as:

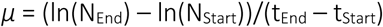

where N is the estimated cell abundance at the experimental Start and End. Bacterial mortality was estimated as:

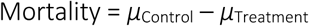

where *μ*_Control_ is the average bacterial growth rate in the controls and *μ*_Treatment_ is the bacterial growth rate in the corresponding treatment.

### DNA extraction, Illumina amplicon sequencing and bioinformatics

Total community DNA was extracted from the filters using the DNeasy PowerSoil Kit (QIAGEN, Hilden, Germany) according to the manufacturer’s instructions with minor modifications in the lysis step; mechanical lysis was achieved by bead-beating pre-cut filters in a Retsch MM2 swing mill (Retsch GmbH, Haan, Germany) for 5 min at a rate of 70 strokes per second. Extracted DNA was amplified with primer pairs targeting the V4 region of the 16S rRNA gene (515f: 50-GTGYC AGCMGCCGCGGTAA-30, 806r: 50-GGACTGCNVGGGTWTC TAAT-30 (37), coupled to custom adaptor-barcode constructs. PCR amplification and Illumina MiSeq library preparation were carried out by LGC Genomics (Berlin, Germany). Adaptor and primer clipped sequences were processed and denoised using the DADA2 pipeline (v.1.2.0 (38) using R (v.4.1.2 (39)). All forward and reverse Illumina reads were simultaneously trimmed to 200 bp, and filtered out if not meeting the quality threshold (maxEE=2, minLen=175). The filtered sequences were then de-replicated and error rates were used to infer sample sequences into amplicon sequence variants (ASVs). Paired forward and reverse sequence reads were merged and followed by a chimera sequence check and removal. Sequences have been submitted to the NCBI short read archive (bioproject number and accession number will be added here). The resulting ASVs were used to construct a table containing relative abundances of ASVs across all samples. The Silva search and classify function in combination with the Silva database (v.138.1, (40) was used to classify ASVs to taxa at the lowest taxonomical levels possible. Further, ASVs assigned to chloroplast or mitochondria and ASVs read counts contributing less than 0.01% to the overall library size were excluded. Background bacteria ASVs were defined as those ASVs present in the respective protist samples (protist culture combined with prey bacteria) but not in the control samples (prey only) at the experimental start. As the scope of this study is understanding the impact of the selected protist on different bacterial communities originating from natural lake communities, we excluded background bacteria ASVs from the dataset by subsequently subtracting them from all samples. 16S rRNA gene copy number (GCN) was estimated as a functional trait proxy for bacterial growth strategy, assuming that bacterial taxa with high 16S rRNA GCN have a higher potential growth rate owing to rapid expression of ribosomes (41,42). GCN was estimated per ASV based on the rrnDB (v.5.7, (43) and standardized per sample by multiplying the GCN belonging to an ASV with its relative contribution to library size.

### Statistical analyses

All statistical analyses were performed in R (v.4.1.2 (39)). Figures were generated using the ggplot2 package (v.3.3.6 (44). ASV richness was calculated by rarefying the whole dataset read counts to the lowest number of reads in a sample. Pielou’s evenness was calculated as H/log(rarefied richness), where H is the Shannon diversity index (45). Spearman’s rank correlation analysis was used to assess the relationship between protist growth, bacterial mortality, bacterial community turnover, changes in GCN, richness and evenness. The change in GCN, richness and evenness ware calculated by subtracting the start value from the end value of the parameter in question. Significant differences between treatments were evaluated per experiment by one-way ANOVA followed by Tukey-HSD post-hoc test. Multivariate statistical analyses were carried out using the vegan package (v.2.6.2 (46)). Non-metric multidimensional scaling was performed on Hellinger-transformed sequence counts using Bray–Curtis dissimilarity to visualize similarities in ASV composition between samples. PERMANOVA was used to evaluate variation in community composition in response to lake and treatment. Differential abundance analyses were used as implemented in the DEseq2 R package (v.1.36.0 (47) to identify differently abundant ASVs between control and protist treatments per lake using raw sequence counts of all ASVs (i.e. without excluding ASVs).

## Results

### Bacterial community composition

Illumina amplicon sequencing of 16S rRNA gene fragments resulted in 8.57 million 16S rRNA gene reads with an average of 64 957 reads per sample (min =18 882, max = 183 337). In total, 1130 unique sequence reads where identified, of which 316 could be assigned to the ‘Background bacteria’ (bacteria pre-existing in the non-axenic protist culture), while the rest were associated with the lake communities. As the focus of this study is set on the impact on enriched lake communities, sequences belonging to the background bacteria, protist chloroplasts and mitochondria were excluded from most data analyses, resulting in 796 ASVs. Further, we refer to the ASVs as bacteria, due their overwhelming abundance compared to Archaea (98.92% of ASVs and 99.19% of reads).

Overall, the prey bacteria prepared from the different lake inocula displayed a community composition representative of freshwater systems, with dominant members belonging to common freshwater genera within the orders *Burkholderiales, Enterobacterales* and *Pseudomonadales* (*Gammaproteobacteria*, Figure 1). *Bacteroidota* were also abundant in almost all lakes, except Obersee. Some lake prey communities showed distinctive features with certain bacterial groups being abundant and not shared between all lakes. *Alphaproteobacteria* and *Pseudomonadales* were more abundant in Erlaufsee, Klostersee and Lunzer See, *Bdellovibrionota* in Erlaufsee and Lunzer See, *Enterobacterales* in Klostersee and *Actinobacteria* were more abundant in Hubertussee than in the remaining lakes. These differences resulted in a strong separation of start prey communities by lake origin (PERMANOVA R^2^=0.95, p<0.001) while no grouping per protist treatment was detectable (PERMANOVA R^2^=0.05, p=n.s., Figure 2a). The same separation manifests no matter if background bacteria are included or excluded from the NMDS calculations (for comparison see also Figure S1). Mittersee and Chiemsee start communities showed an overlap, but NMDS analysis of bacterial start communities from only these two lakes showed a clear separation per lake (not shown). Thus, the bacterial prey communities varied depending on lake of origin, with their overall composition reflecting the diversity of natural bacterial communities. After the 48h incubation lake origin still had a strong influence on the bacterial communities (PERMANOVA R^2^=0.80, p<0.001), but now the influence of protists was also significant (PERMANOVA R^2^=0.11, p<0.001, Figure 2b).

**Figure 1.**
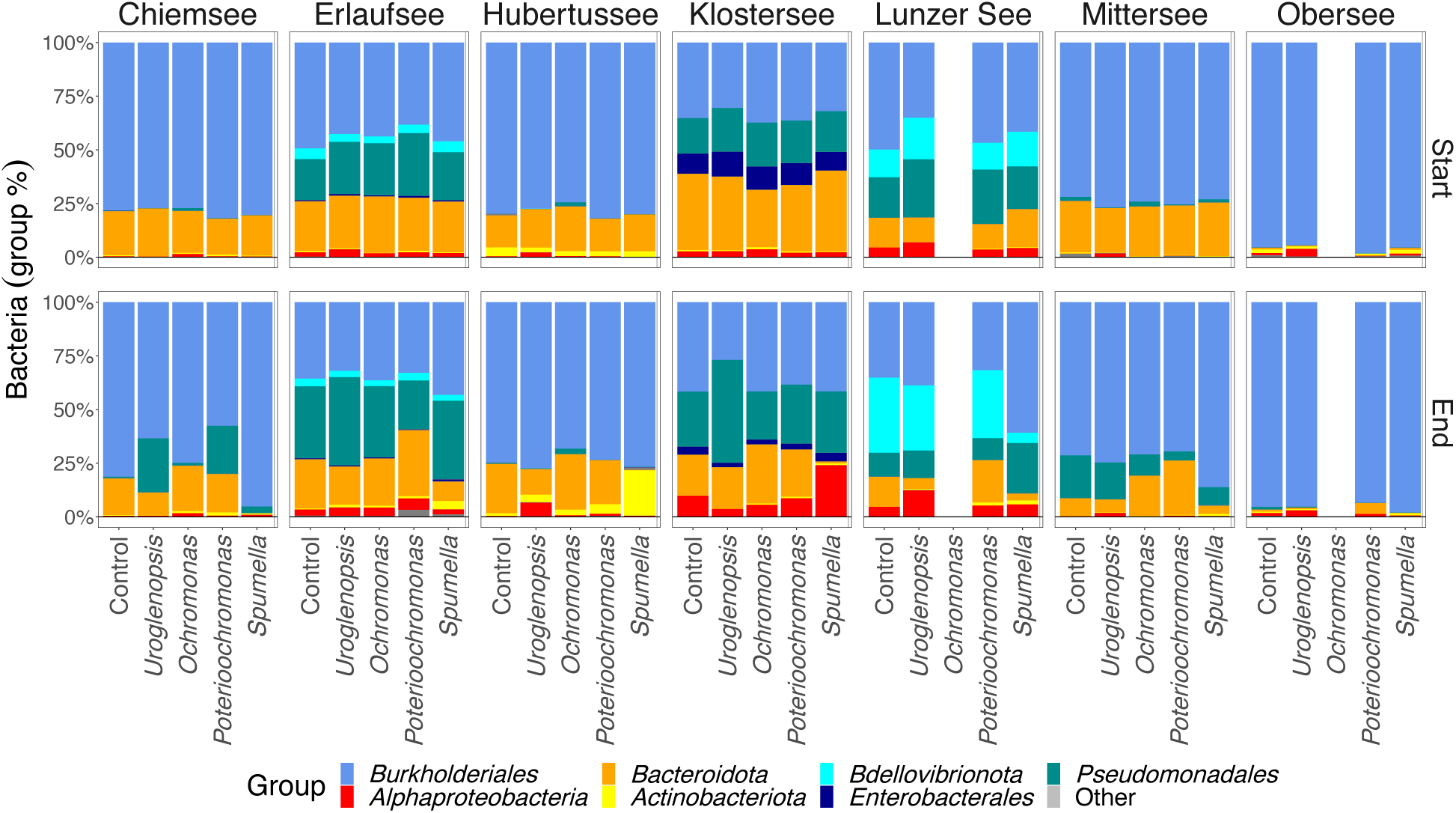
Added prey communities showed distinctive features on a broad taxonomic level depending on lake of origin. Relative abundance of bacterial groups in all treatments per experiment at the start and end of the 2-day experimental incubation. At the experimental end the mean abundance of triplicate samples per treatment and lake are presented. Error bars are not shown. Bacterial groups are presented at the Phylum level, except *Proteobacteria* which are split up to the class *Alphaproteobacteria* and *Gammaproteobacteria* orders (*Burkholderiales, Enterobacterales* and *Pseudomonadales*). Other includes diverse groups present with less than 1% abundance.

**Figure 2.**
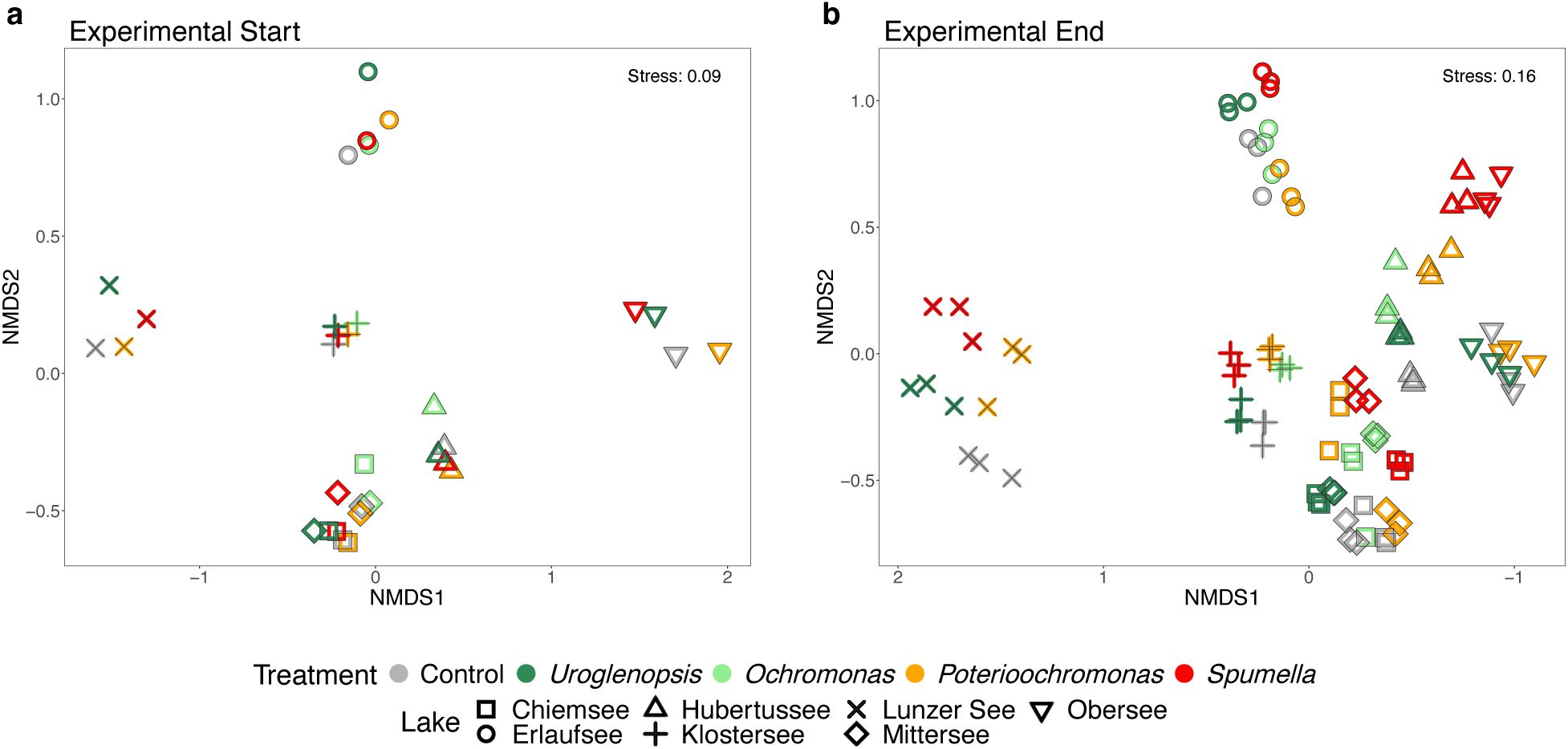
Bacterial prey communities were separated by lake of origin, yet protist impact was evident after 48h. Added bacterial communities present at the start (a) and end (b) of all experimental incubations derived from NMDS-ordinations based on Bray–Curtis dissimilarities.

### Bacterial community composition changes were related to protist growth and grazing rates

Growth and grazing rates and induced changes in bacterial community composition (BCC) increased towards pure heterotrophic nutrition. Bray-Curtis dissimilarity between start and end samples (Figure 3a, see Figure S2 for the matching NMDS plot) increased significantly in most *Spumella* and *Poteriochromonas* incubations. Contrary, *Uroglenopsis* and *Ochromonas* in some experiments did not induce a change in Bray-Curtis dissimilarity (as well *Poteriochromonas* in Erlaufsee), with BCC changing more only in three incubations with *Uroglenopsis*, while in most *Ochromonas* treatments BCC changed significantly less than in the matching control (Figure 3a).

**Figure 3.**
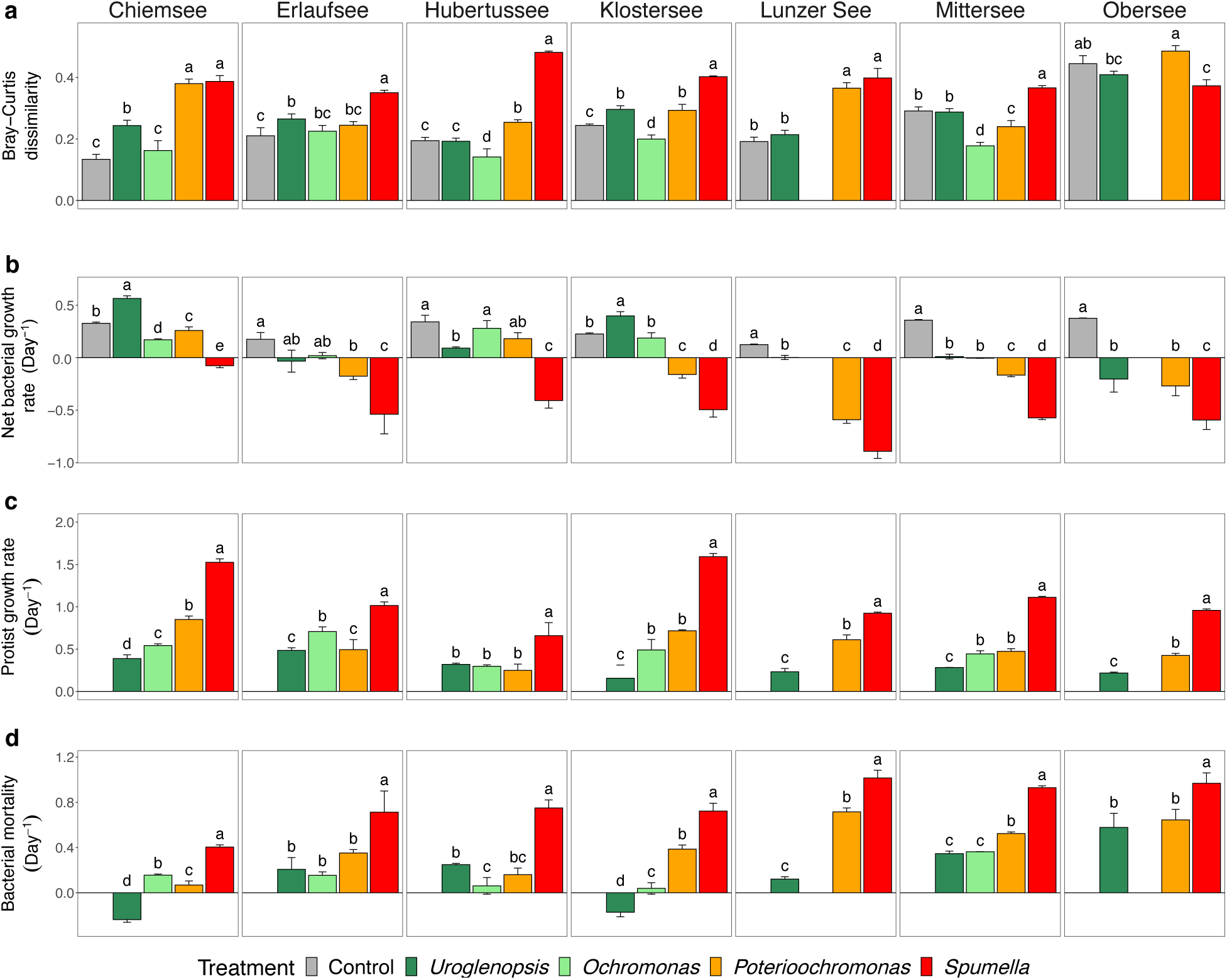
Shift in bacterial community composition between start and end samples (Bray-Curtis dissimilarity, a), bacterial net growth rate (b), protist growth rate (c) and bacterial mortality (d) per treatment and lake. Values are means of triplicates; error bars represent SD. Different letters indicate significant differences between the treatments per lake incubation (one-way ANOVA followed by post hoc Tukey test).

The presence of the protists induced changes in bacterial net growth in almost all incubations (Figure 3b). Most protist incubations induced lower bacterial growth, most pronounced in *Spumella* and *Poteriochromonas* treatments. In contrast, Chiemsee and Klostersee *Uroglenopsis* supported significantly higher bacterial net growth, without any impact in Erlaufsee and in the remaining experiments it significantly reduced bacterial net growth. *Ochromonas* significantly reduced bacterial growth only in Chiemsee and Mittersee. The protists significantly differed in their own growth rates and induced bacterial mortality, with *Spumella* exhibiting the highest growth and ingestion rates and *Uroglenopsis* and/or *Ochromonas* overall the lowest (Figure 3c, d).

The predicted gradient from photo- to heterotrophy (*Uroglenopsis* > *Ochromonas* > *Poteriochromonas* > *Spumella)* is well illustrated through the positive relationship between protist growth and bacterial mortality, and the matching histograms of these parameters (R^2^=0.537, p<0.001, Figure 4). Bray-Curtis dissimilarity between start and end samples exhibited a significant positive relationship with protist growth (R^2^=0.422, p<0.001) and bacterial mortality (R^2^=0.641, p<0.001). In addition to these overall trends, for a given protist alone, only *Ochromonas* growth showed a positive relationship with bacterial community turnover (R^2^=0.757, p<0.01), while *Spumella* growth rates and bacterial mortality had a negative correlation (R^2^=-0.457, p<0.05).

**Figure 4.**
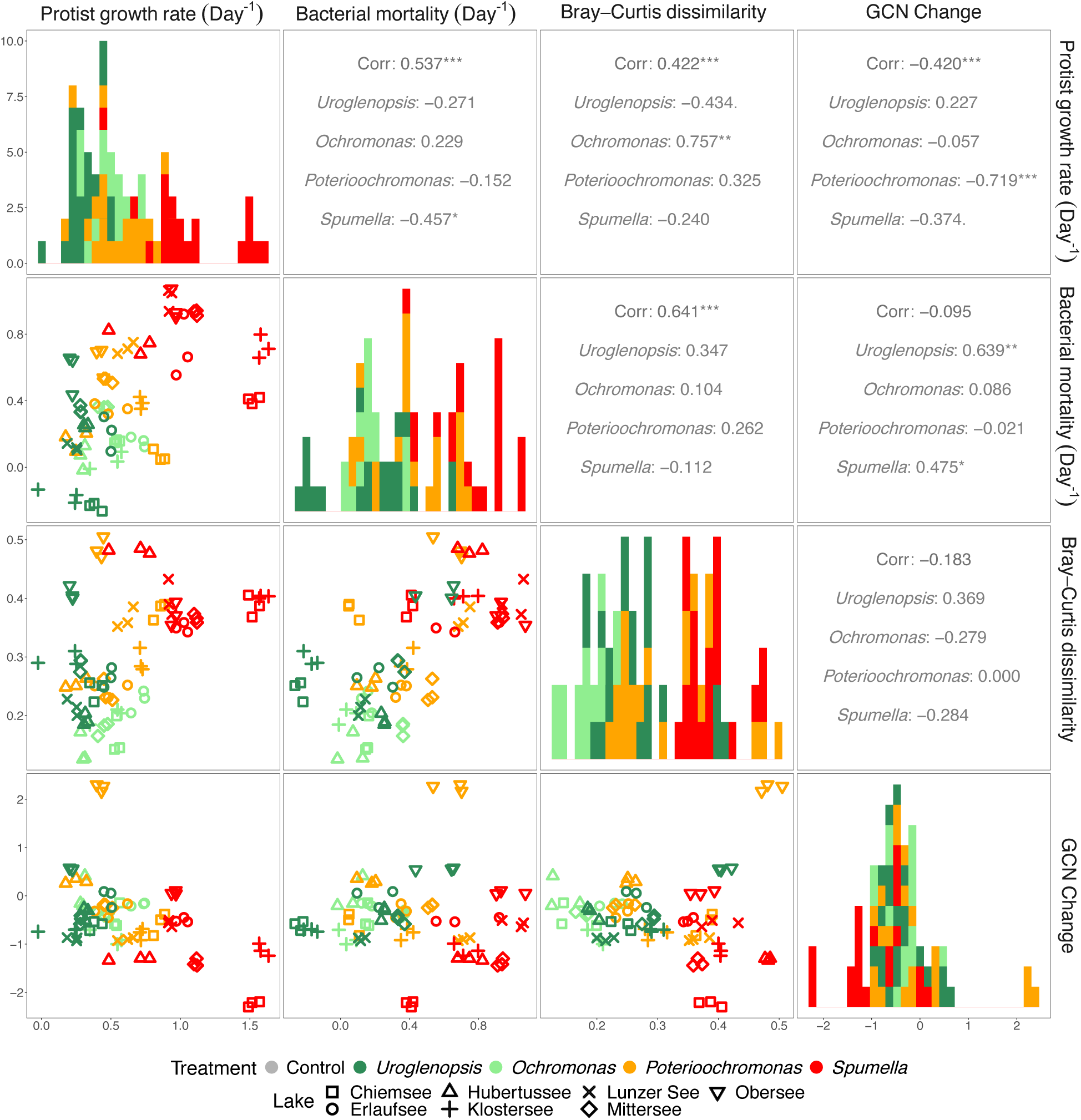
The relationship between protist growth, bacterial mortality, shift in bacterial community composition between start and end samples (expressed as Bray-Curtis dissimilarity) and change in average 16S rRNA gene copy number during the experimental incubations (GCN change). Spearman’s rank correlation, R^2^ values are indicated by numbers, and p values by .(p<0.1), *(p<0.05), ** (p<0.01), *** (p<0.001).

### Impact of protists on bacterial prey traits and taxonomic identity

The copy number of the 16S rRNA gene, as a proxy for bacterial growth (41,42), could be estimated on 59.05% of the bacterial ASVs. This trait varied among the prey bacterial communities, and a gradient of average estimated 16S rRNA gene copy number (GCN) was present among the communities depending on lake (Figure S3a), with the lowest GCN in Obersee and highest in Erlaufsee. Incubations with *Spumella* which featured the highest protist growth rates and bacterial mortality also led to the biggest decrease in GCN in all experiments except in Lunzer See (Figure 4, S3b). Overall, protist growth rate had a negative relationship with GCN change (R^2^=-0.420, p<0.001), with higher protist growth rates decreasing average prey GCN. *Poteriochromonas* (R^2^=-0.719, p<0.001) and *Spumella* (R^2^=-0.374, p<0.1) growth rates were as well negatively related to GCN change. There was no relationship between overall mortality and GCN change, but for a given protist alone, i.e. in *Spumella* (R^2^=0.475, p<0.05) and *Uroglenopsis* (R^2^=0.639, p<0.01) treatments increasing prey mortality related to a GCN increase. Growth and mortality rates in *Ochromonas* incubations did not show any relation to GCN changes.

Bacterial mortality correlated positively with the change in richness in *Uroglenopsis* (R^2^=0.464, p<0.05, Figure 5) and *Ochromonas* (R^2^=0.507, p<0.1) and with the change in evenness in *Uroglenopsis* (R^2^=0.423, p<0.1), *Ochromonas* (R^2^=0.604, p<0.05). Contrary, a negative relationship with richness and evenness change formed in *Poteriochromonas* and *Spumella* (p=n.s.). Overall the change in bacterial richness and evenness had a positive relationship (R^2^=0.444, p<0.001), with R^2^ decreasing from primarily phototrophic towards primarily phagotrophic protist (*Uroglenopsis*: R^2^=0.457, p<0.01, *Ochromonas:* R^2^=0.493, p<0.1, *Poterioochromonas*: R^2^=0.384, p<0.1, *Spumella*: R^2^=0.078, p=n.s.).

**Figure 5.**
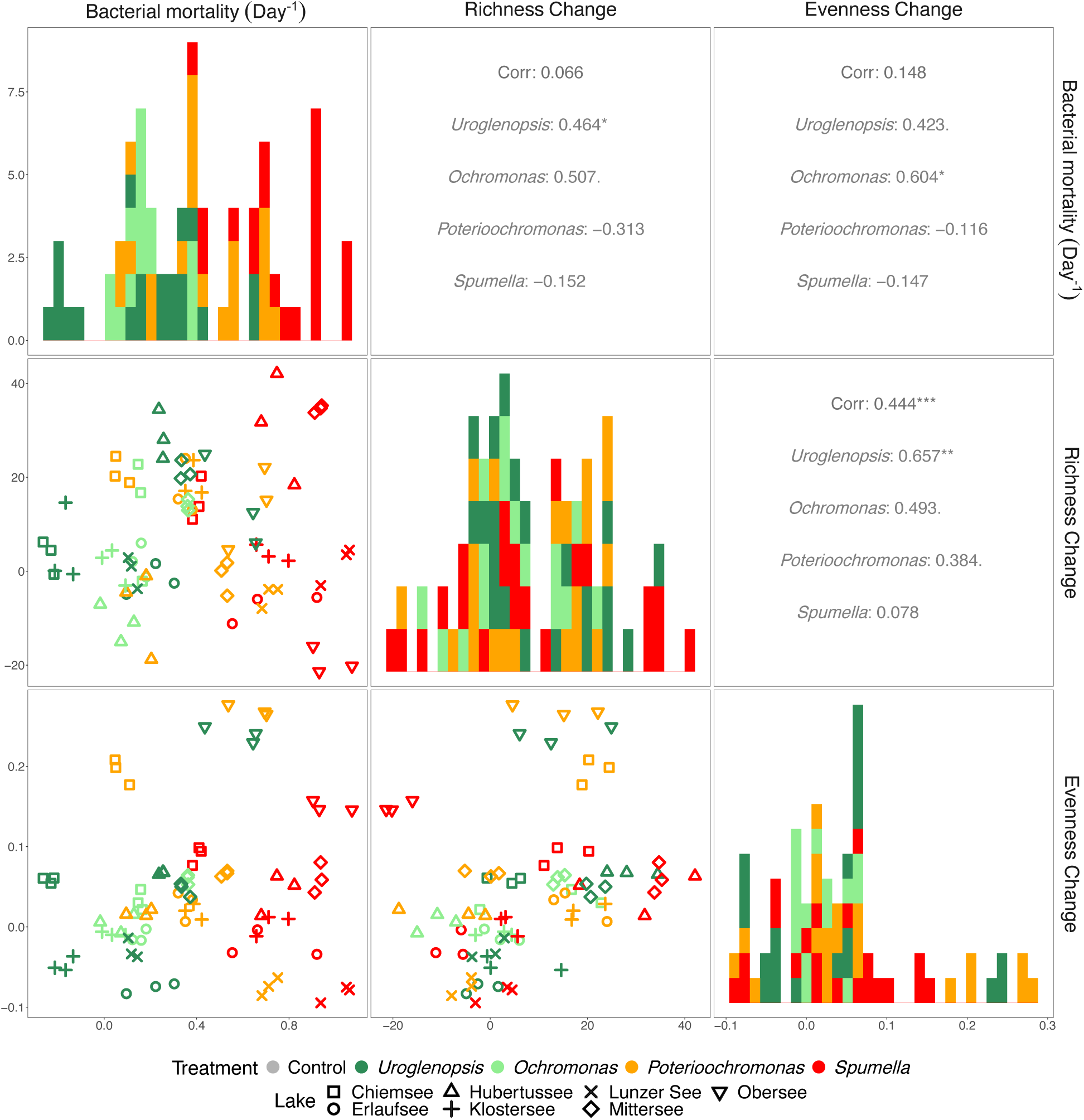
The relationship of bacterial mortality with the change in bacterial species richness (Richness change) and evenness (Evenness change) during the experimental incubations. Spearman’s rank correlation, R^2^ values are indicated by numbers, and p values by .(p<0.1), *(p<0.05), ** (p<0.01), *** (p<0.001).

General patterns independent of lake origin could be observed through the estimation of significantly differing ASVs between protist incubations and their matching control per lake (Figure 6, p<0.01, DEseq2 analyses). The overall ASV changes coincide with a decrease of abundant and high GCN ASVs towards clear phagotrophic nutrition. For more details about the exact change of ASVs related to their abundance and GCN see Table S2 and S3. Strikingly, all protist incubations led to a significant increase of *Actinobacteria* ASVs. Prominent was as well the significant decrease of *Bacteroidota* ASVs in all *Spumella* samples. On the other hand, *Bacteroidota* had a varying response in incubations with the mixotrophs, for example mainly increasing in *Ochromonas* incubations. *Alphaproteobacteria* and *Gammaproteobacteria* also showed a varying response differing between protist, with *Alphaproteobacteria* increasing in *Ochromonas* and *Poterioochromonas* and *Gammaproteobacteria increasing in Uroglenopsis*. Generally, *Proteobacteria* decreased towards strict heterotrophic treatments. The few remaining significantly different ASVs belonged to *Verrucomicrobiota, Bdellovibrionota, Campylobacterota, Chloroflexi* and *Patescibacteria* (for more details about the exact change of ASVs per bacterial group see Table S4).

**Figure 6.**
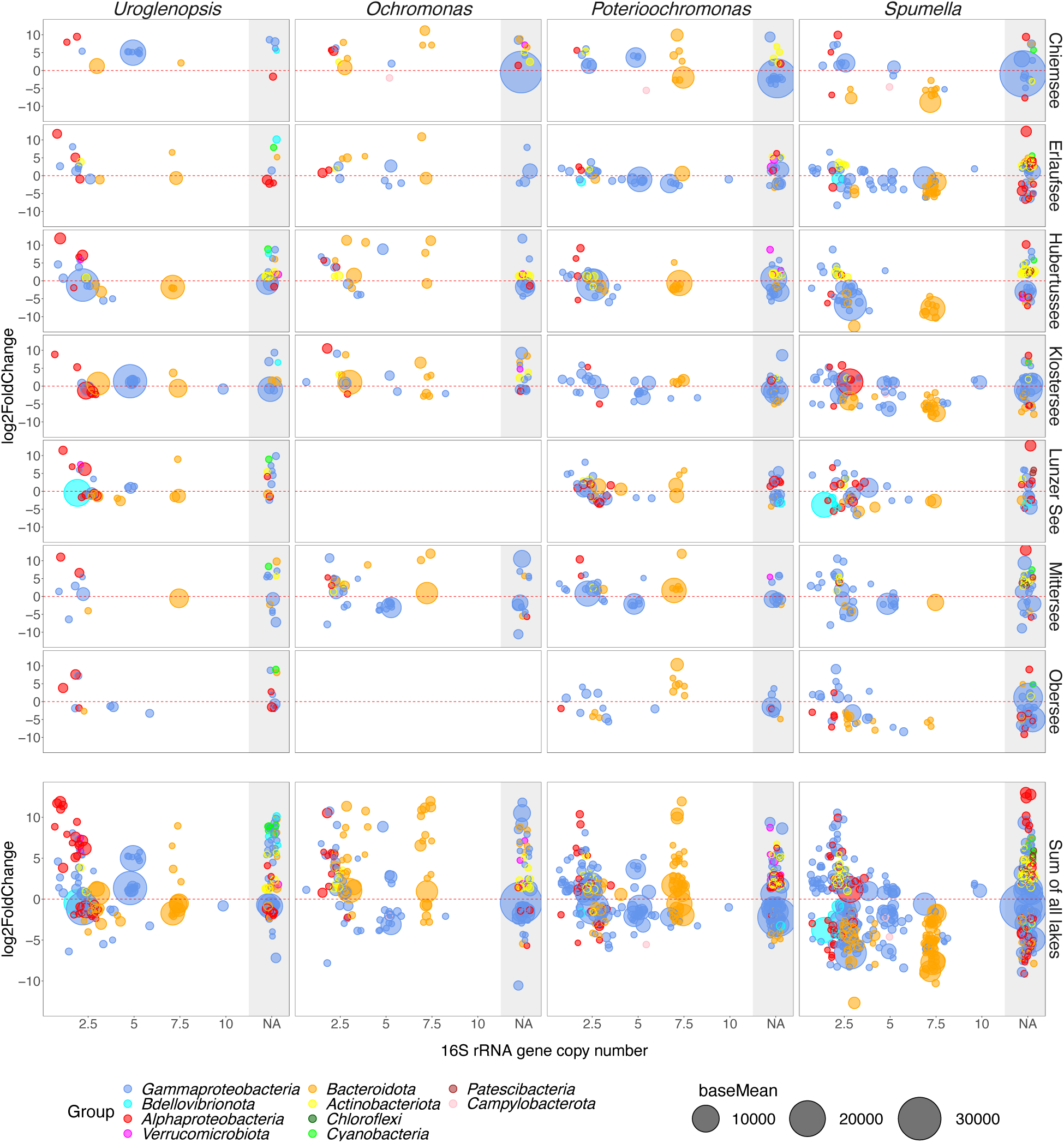
Lake specific bacterial ASVs significantly differing between the control and protist treatment at the end of incubation per protist and separated per lake and as sum of lake experiments. Each circle represents an individual ASV. ASVs above the dotted red line are significantly more present in the protist in comparison to the control, and below significantly less present (presumably removed via grazing). The position of each circle on the x-axis of is proportional to the rRNA Copy number of that ASV, while the area of each circle is proportional to the abundance of an ASV (baseMean, across experiment).

## Discussion

### Protist grazing affected bacterial communities depending on their trophic mode

The observed bacterial community changes supported our overall hypothesis that a protist nutritional mode predicts its impact on bacterial communities. Bacterial community composition (BCC) changes were accompanied by increasing protist growth and bacterial mortality along the gradient from phototrophy towards heterotrophy. Bacterial community shifts were consistent with structuring by ingestion, illustrated by the robust positive relationship between BCC change and bacterial mortality, although effects on BCC via other mechanisms cannot be completely ruled out. The clear heterotroph *Spumella* and the primarily phagotrophic facultative mixotrophs *Poterioochromonas* induced stronger changes in BCC, while the primarily phototrophic obligate mixotrophs (OM) *Uroglenopsis* and *Ochromonas* exhibited a weaker impact on BCC, which in some experimental incubations resulted in a stabilizing effect on BCC. However, the positively correlated growth of *Ochromonas* with BCC changes implies that OM might lose their stabilizing effect after reaching a certain abundance. This can be related to the fact that the growth and survival of heterotrophic protists completely depends on bacterial ingestion, while mixotrophs ingest less bacteria per cell division the more they use phototrophic nutrition (16). The mixotrophic dual strategy creates the potential for interactions with bacteria beyond predation. Perhaps mixotrophs might achieve this stabilizing effect by cultivating the bacteria they graze through exudates. Facilitation of bacterial growth through organic exudates is known for phototrophic algae (48,49). Such interactions between mixotrophic protists and bacteria are not well studied, but it is known that nutrient release of mixotrophic protists depends on the protist in question, present prey, nutrient and light levels (18,20,50).

### Protist impact on bacterial prey communities was related to bacterial 16S rRNA gene copy number

16S rRNA gene copy number (GCN) is a bacterial trait linked to maximum growth rate (41,42) that can be predicted based on the taxonomic composition of 16S rRNA-based sequencing datasets. We found that the average GCN of a given prey community was affected differently per protist. The heterotrophic protist *Spumella* induced the highest GCN decrease in all experiments, suggesting selective grazing by *Spumella* on fast-growing bacteria. Fast-growing bacteria are presumably abundant and frequently encountered, but often lack protective mechanisms and apply high growth rates to escape grazing pressure (31). Ingestion of bacteria with such characteristics would not require active protist selection. On the other hand, high GCN may also correlate with high ribosomal content, which could translate into better food quality. We did not measure prey stoichiometry, but high and low GCN bacteria might differ in their stoichiometry, with fast growing metabolically active bacteria being more P rich (25). The ideal prey of a heterotrophic consumer has a higher C:(N,P) than the consumer (51). However, heterotrophic protists as fast growing specialist might prefer P-rich prey, due increased demands for ribonucleic acids (RNA), and thus P at high growth rates (25,51). Strictly light-dependent mixotrophs such as *Uroglenopsis* gain most of their energy and carbon through photosynthesis, while using prey primarily as N and P source. Thus, potential selectivity of mixotrophs related to prey C:N:P content would depend on present nutrient and light levels.

### Protist nutritional strategy exerted a consistent influence on bacterial prey on a broad taxonomical level

The protists induced pronounced changes in bacterial groups across all experiments, pointing towards potential general selectivity patterns applying already at higher bacterial taxonomical levels. *Actinobacteria* always increased significantly in comparison to the controls regardless of protist treatment, indicating some type of general grazing resistance. Our results thus reinforce similar observations of *Actinobacteria* that have been made in several other studies focusing on bacterivorous protists (example:(52–55). Apparently, it is almost a rule that the presence of bacterivorous protist creates an advantage for *Actinobacteria* over other bacteria. So far, formation of aggregates and their small cell size are suggested to be predation defense mechanism, implying a size ratio mismatch for most bacterivores (32,52). This in combination with their cell walls characteristics and possible aggravated digestion of them as gram positive bacteria could lead to their general avoidance by phagotrophic protists (56,57).

Most bacteria belonging to *Alphaproteobacteria, Gammaproteobacteria* and *Bacteroidota* showed a varying response depending on protist treatment, but followed a gradual change in the ratio of a significant increase-decrease matching to the nutritional strategy of the protist, with a stronger decrease of these groups towards the heterotrophic end. Interestingly, *Spumella* selectively depleted *Bacteroidota* ASVs in our experiments, indicating a preference for this taxonomical group as prey. Previous studies confirmed initial strong selection of heterotrophic protist for *Bacteroidota*, which shifted towards avoidance in parallel with bacterial cell-size increase and formation of grazing resistant morphotypes (53,58,59). This is in accordance to the expectation that preferred bacteria get less available or develop defense mechanisms with time (32).

Other studies found increasing preference for certain *Betaproteobacteria* subgroups with time (53,54). However, in our case *Alpha*- and *Beta-proteobacteria* did not show a clear trend. This might be related to our short incubation time were a preference shift from *Bacteroidota* to *Beta-proteobacteria* could not be noticed. While P*roteobacteria* ASVs generally decreased towards clear heterotrophy, *Ochromonas* and *Poterioochromonas* mainly supported *Alphaproteobacteria* growth. In addition, *Ochromonas* as well supported *Bacteroidota* growth. *Alphaproteobacteria* are able to utilize macroalgal polysaccharides and are often found to be positively associated with phototrophic protist (60). Potentially, these bacterial groups could have an advantage from interactions with mixotrophic protist by profiting from algal exudates, explaining their higher abundance only in mixotrophic treatments.

### Implications of different protist nutritional strategies under current and future climate conditions

The impact intensity on bacterial communities increased towards clear heterotrophic nutrition, with OM potentially having a stabilizing impact on bacterial communities. The identity of the prevailing protist directly affects BCC, suggesting that OM are more flexible in terms of ingested prey. Strong selectivity in *Spumella* and *Poteriochromonas* was as well already confirmed in other studies (61,62). It should be tested if this trend holds true for other taxonomic lineages (especially outside Chrysophytes) and if this is linked to metabolomics flexibility as arising from their dual strategy. Selectivity in marine dinoflagellates was found to increase towards heterotrophic nutrition (63). The study suggested that primarily phototrophic mixotrophic dinoflagellates trade fast growth for feeding on diverse prey, to gain an advantage over resource limited specialists. However, in our study mixotrophic selective grazing might had not been detected due to their low grazing rates in combination with the used methods. Our approach relies on the overall change in BCC created by the protist presence, but does not allow to distinguish alternative interactions that might lead to similar outcomes, e.g. avoidance of prey vs. compensatory growth.

Primarily heterotrophic mixotrophs and strictly heterotrophic protists likely dominate bacterivory, while light-dependent mixotrophs seem quantitatively less important. In natural systems, we overall see an inverse relationship between the trophic state of a lake and share of mixotrophs in the phytoplankton (64). The dual mode of nutrition seen in mixotrophy comes with extra costs and is evident from overall lower maximum growth rates in mixo-vs. heterotrophic consumers (65,66). Nevertheless, in many systems mixotrophs dominate bacterivory (11,67). Especially in environments with low bacterial abundance and saturating light the dual strategy allows mixotrophs to outcompete heterotrophic protists for their shared prey (65,68). In systems where mixotrophs bloom they can play an important role in shaping bacterial communities, although individual grazing rates may be low compared to heterotrophic protists. Interestingly, the *Uroglenopsis* species used in this study is a mixotroph relying more on photosynthesis, with overall low growth and grazing rates, yet the same species is regularly forming summer blooms in its originating lake, the Lunzer See (own observation).

Future climate change scenarios predict an increase in extreme weather events, leading to increased physical stress, reduced light intensities and possibly elevated dissolved organic matter levels of aquatic systems (69–71). This combined with temperature rising (72) might favor specialist over generalist, and heterotrophic over phototrophic metabolism in protists (73). In addition warmer temperatures may case elongated lake stratification, leading to nutrient decline (74), and favoring OM in illuminated lake layers. Our study suggests their dominance would have a stabilizing effect on BCC, until they reach a certain abundance. Thereby, any change supporting the dominance of more heterotrophic protist and prolonging blooms of OM would lead to higher bacterial consumption and dominance of grazing protected bacteria linked to prey selectivity specific for the protist group in question. Such differing structuring impacts on bacterial communities might consequently affect the overall food web and element cycling. Anyway, we hypothesize that even blooming OM have a buffering effect on BCC if set in comparison to the impact of heterotrophic bacterivores.

This study is the first to tackle the impact of protist bacterial community structure with different trophic modes, covering a gradient from primarily phototrophic to strictly heterotrophic bacterivorous protists. We highly encourage future studies to consider different nutritional strategies within protists when addressing protist-bacteria interactions.

## Acknowledgements

This research was funded by the DACH project ‘The impact of mixotrophic bacterivory on microbial food webs in changing lake ecosystems (LakeMix)’, Austrian Science Fund (FWF) I 3311-B25 and German Research Foundation (DFG) BE 6194/1-1. M. IvankoviĆ acknowledges further support by the Austrian Agency for International Cooperation in Education and Research (OeAD-GmbH) through the Marietta Blau scholarship, and support from the European Union’s Horizon 2020 research and innovation programme under grant agreement 871081. The authors thank Christian Preiler and Stephanie Grubner for practical help during the experimental phase.

## Data availability statement

The materials described in this manuscript, including all relevant raw data, will be freely available to any researcher wishing to use them for non-commercial purposes, without breaching participant confidentiality.

