## Supplementary material for "Top-down structuring of freshwater bacterial communities by mixotrophic and heterotrophic protists"

Supplement

**Supplementary Table 1** Lake characteristics.

| Lake | Trophic status | Country | Coordinates |
|---|---|---|---|
| Chiemsee | Mesotrophic | Germany | 47°52'11.61"N, 12°27'12.58"E |
| Erlaufsee | Oligotrophic | Austria | 47°47'33.42"N, 15°16'19.62"E |
| Hubertussee | Mesotrophic | Austria | 47°48'28.86"N, 15°21'59.01"E |
| Klostersee | Eutrophic | Germany | 47°58'28.32"N, 12°27'8.49"E |
| Lunzer See | Oligotrophic | Austria | 47°51'13.43"N, 15° 3'5.66"E |
| Mittersee | Oligotrophic | Austria | 47°49'38.09"N, 15° 4'33.04"E |
| Obersee | Oligotrophic | Austria | 47°48'16.56"N, 15° 4'27.56"E |

**Supplementary Table 2** High abundant ASVs (below average baseMean) and low abundant ASVs (above average baseMean) significantly differing between the Control and protist treatment at the end of incubation. Shown are ASVs of lake experiments together, which significantly increased (+), decreased (-) and the log response ratio (log(+/-)) between increased and decreased bacterial ASVs per protist. Average baseMean of lake experiments = 553.57.

| | *Uroglenopsis americana* | | | *Ochromonas* cf. *perlata* | | | *Poterioochromonas malhamensis* | | | *Spumella* sp. | | |
|---|---|---|---|---|---|---|---|---|---|---|---|---|
| | + | - | log(+/-) | + | - | log(+/-) | + | - | log(+/-) | + | - | log(+/-) |
| baseMean < 553.57 | 98 | 50 | 0.29 | 85 | 46 | 0.27 | 136 | 121 | 0.05 | 199 | 263 | -0.12 |
| baseMean > 553.57 | 11 | 18 | -0.21 | 13 | 7 | 0.27 | 21 | 26 | -0.09 | 20 | 41 | -0.31 |

**Supplementary Table 3** High GCN ASVs (< average GCN) and low GCN ASVs (> average GCN) significantly differing between the Control and protist treatment at the end of incubation. Shown are ASVs of lake experiments together, which significantly increased (+), decreased (-) and the log response ratio (log(+/-)) between increased and decreased bacterial ASVs per protist. Average GCN of lake experiments = 3.79.

| | *Uroglenopsis americana* | | | *Ochromonas* cf. *perlata* | | | *Poterioochromonas malhamensis* | | | *Spumella* sp. | | |
|---|---|---|---|---|---|---|---|---|---|---|---|---|
| | + | - | log(+/-) | + | - | log(+/-) | + | - | log(+/-) | + | - | log(+/-) |
| GCN < 3.79 | 41 | 34 | 0.08 | 46 | 9 | 0.71 | 72 | 39 | 0.27 | 93 | 99 | -0.03 |
| GCN > 3.79 | 22 | 12 | 0.26 | 20 | 20 | 0.00 | 34 | 42 | -0.09 | 16 | 101 | -0.80 |

**Supplementary Table 4** Bacterial ASVs significantly differing between the Control and protist treatment at the end of incubation. Shown are ASVs of lake experiments together, which significantly increased (+), decreased (-) and the log response ratio (log(+/-)) between increased and decreased bacterial ASVs per protist.

| | *Uroglenopsis americana* | | | *Ochromonas* cf. *perlata* | | | *Poterioochromonas malhamensis* | | | *Spumella* sp. | | |
|---|---|---|---|---|---|---|---|---|---|---|---|---|
| | + | - | log(+/-) | + | - | log(+/-) | + | - | log(+/-) | + | - | log(+/-) |
| *Actinobacteria* | 8 | 0 | 8/0 | 16 | 0 | 16/0 | 20 | 1 | 1.30 | 41 | 1 | 1.61 |
| *Alphaproteobacteria* | 18 | 19 | -0.02 | 11 | 4 | 0.44 | 21 | 9 | 0.37 | 29 | 36 | -0.09 |
| *Bacteroidota* | 14 | 20 | -0.15 | 33 | 7 | 0.67 | 30 | 29 | 0.01 | 6 | 105 | -1.24 |
| *Bdellovibrionota* | 4 | 1 | 0.60 | 0 | 0 | - | 0 | 2 | -∞ | 0 | 6 | -∞ |
| *Campylobacterota* | 0 | 0 | - | 0 | 1 | -∞ | 0 | 1 | -∞ | 0 | 2 | -∞ |
| *Chloroflexi* | 0 | 0 | - | 0 | 0 | - | 0 | 0 | - | 2 | 0 | 2/0 |
| *Cyanobacteria* | 5 | 0 | 5/0 | 0 | 0 | - | 0 | 0 | - | 6 | 0 | 6/0 |
| *Gammaproteobacteria* | 57 | 28 | 0.31 | 35 | 41 | -0.07 | 80 | 105 | -0.12 | 125 | 153 | -0.09 |
| *Patescibacteria* | 0 | 0 | - | 0 | 0 | - | 0 | 0 | - | 9 | 0 | 9/0 |
| *Verrucomicrobiota* | 3 | 0 | 3/0 | 3 | 0 | 3/0 | 6 | 0 | 6/0 | 1 | 1 | 0 |
| Total | 109 | 68 | 0.20 | 98 | 53 | 0.27 | 157 | 147 | 0.03 | 219 | 304 | -0.14 |

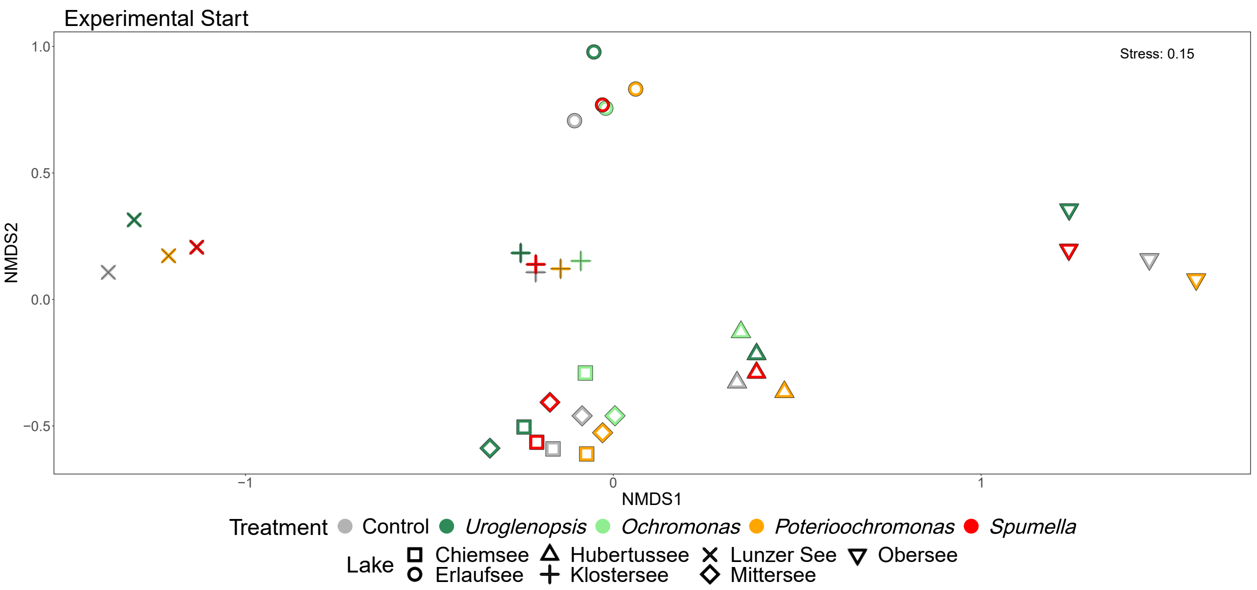


**Supplementary Figure 1** Start bacterial prey communities without excluding background bacteria were separated by lake of origin. Background bacteria and added prey communities present at the beginning off all experimental incubations derived from NMDS-ordinations based on Bray–Curtis dissimilarities.

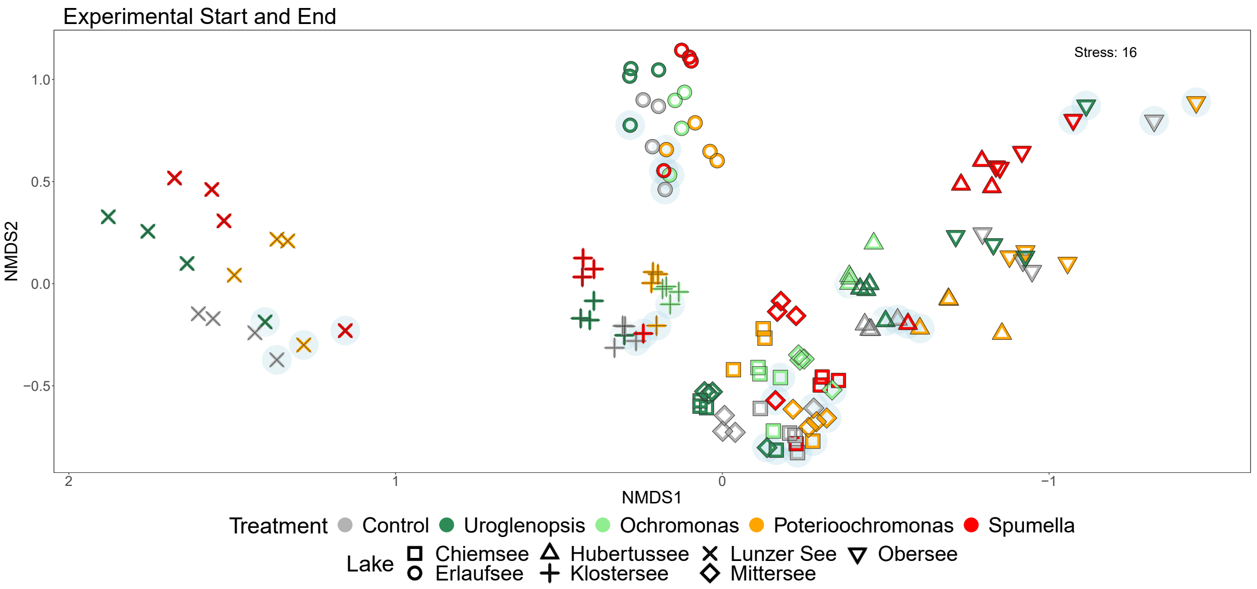


**Supplementary Figure 2** Added bacterial communities present at the beginning and end of all experimental incubations derived from NMDS-ordinations based on Bray–Curtis dissimilarities. Light blue circles indicate samples from the experimental start.

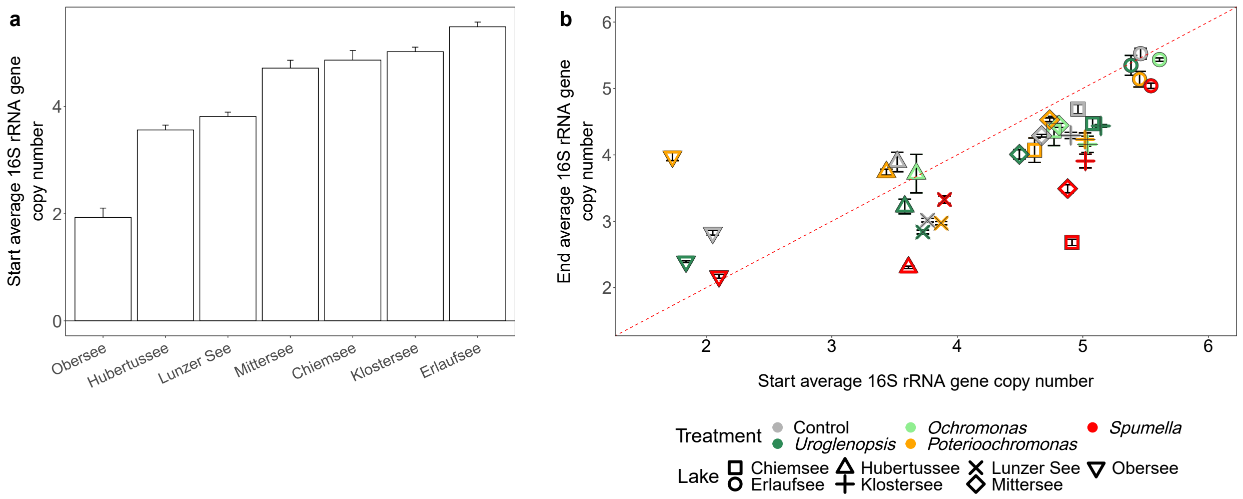


**Supplementary Figure 3** Decrease of bacteria with high 16S rRNA copy numbers was observed for some protists. The average 16S rRNA gene copy number of bacteria at the start of the experimental incubations per lake (a), and the start and end of the experimental incubations per protist and lake (b). Values above the dotted red line increased, and below decreased during the incubations. Error bars represent SD.
